# Environmental selection influences the microbiome of subsurface petroleum reservoirs

**DOI:** 10.1101/2022.09.08.507151

**Authors:** Daniel A. Gittins, Srijak Bhatnagar, Casey R. J. Hubert

## Abstract

Petroleum reservoirs within the deep biosphere are extreme environments inhabited by diverse microbial communities creating biogeochemical hotspots in the subsurface. Despite their ecological and industrial importance, systematic studies of core microbial taxa and associated genomic attributes of the oil reservoir microbiome are limited. This study compiles and compares 343 16S rRNA gene amplicon libraries and 25 shotgun metagenomic libraries from oil reservoirs in different parts of the world. Taxonomic composition varies among reservoirs with different physicochemical characteristics, and with geographic distance. Despite oil reservoirs lacking a taxonomic core microbiome in these datasets, gene-centric metagenomic analysis reveals a functional core featuring carbon acquisition and energy conservation strategies consistent with other deep biosphere environments. Genes for anaerobic hydrocarbon degradation are observed in a subset of the samples and are therefore not considered to represent core biogeochemical functions in oil reservoirs. Metabolic redundancy within the petroleum reservoir microbiome reveals these to be deep biosphere systems poised to respond to changes in redox biogeochemistry. This highlights the potential to use microbial genomics for predicting microbial responses to (bio)engineering perturbations to these subsurface habitats.

## Introduction

The subsurface biosphere is the largest microbial habitat on Earth (Bar-On *et al*. 2018). Microbes inhabiting this environment play an important role in mediating global scale biogeochemical cycling of elements and nutrients (Orcutt *et al*. 2013). Understanding the factors that shape microbial community structure and functional potential helps to uncover these processes and understand how geology and biology interact. The deep biosphere includes diverse terrestrial and marine habitats such as aquifer systems, basaltic ocean crust, buried sediments and petroleum reservoirs. The latter represents an interesting microbial setting in the subsurface context in that energy rich petroleum compounds offer substrates and electron donors, while at the same time hydrocarbons are generally considered toxic to microorganisms (Sikkema *et al*. 1995; Xu *et al*. 2018). Petroleum reservoirs contain an order of magnitude more microorganisms than surrounding sediments at similar depths (Bennett *et al*. 2013), indicating that these systems may be deep biosphere ‘hot spots’ of microbial activity.

The physiology and metabolism of the petroleum reservoir microbiome influences the physicochemical properties of these environments. Crude oil biodegradation changes the composition and physical properties of both liquid and gaseous components of petroleum via the sequential metabolism of hydrocarbons and other compounds (Head *et al*. 2003). Methanogenic oxidation of petroleum hydrocarbons over geological timescales is catalyzed *in situ* by consortia of bacteria and archaea, altering both the chemistry of the remaining oil as well as the levels of CO_2_ and CH_4_ (Jones *et al*. 2008; Milkov, 2011). Interestingly, this oil-altering biogeochemistry appears to only take place in reservoirs that have a burial history that has not included depths hotter than 80–90°C, inactivating these populations by heat sterilization (Wilhelms *et al*. 2001). In situations where sulfate is present, anaerobic respiration of sulfate to sulfide, known as reservoir souring, can be coupled to the oxidation of hydrocarbons directly, or indirectly via oxidation of organic acids or hydrogen derived from crude oil biodegradation (Aitken *et al*. 2013; Gieg *et al*. 2010; Sherry *et al*. 2013). Accurate characterisation of these metabolic processes catalysed by the petroleum reservoir microbiome enables useful predictions of reservoir conditions.

Microbial communities in nature are typically highly complex, comprising thousands of species. A ‘core microbiome’ is considered to be the consistent component of complex microbial assemblages present in a given habitat type, as confirmed by observations of common taxa across several different sampling sites (Turnbaugh *et al*. 2007; Hamady and Knight, 2009; Shade and Handelsman, 2011). Identifying the occurrence of a core microbiome is important for unravelling the ecology of an ecosystem. Commonly occurring microorganisms are likely also important for normal biogeochemical functioning, such that the microbiome helps to define dominant metabolic processes and microbial interactions (Neu *et al*. 2021). Temporal and spatial heterogeneity in microbiota can also occur, resulting in dynamic shifts in functioning across complex microbiomes (Astudillo-García *et al*. 2017). In petroleum reservoirs, identifying common taxa may guide engineering interventions with the potential to manipulate these subsurface communities and the important process they mediate.

The most common method for microbial community profiling in oil reservoirs is to sequence the 16S rRNA gene following PCR amplification. Shotgun metagenome sequencing is now also applied to oil reservoir samples with increasing regularity (Hu *et al*. 2016; Vigneron *et al*. 2017; Christman *et al*. 2020). Metagenomic studies can advance understanding of the microbial ecology of a given ecosystem relative to single-gene amplicon sequencing by revealing information about metabolic genes and functional potential within the sampled community. These techniques are routinely applied in other sectors for biotechnological applications ranging from medicine to bioenergy. Here we provide a meta-analysis of 16S rRNA genes and metabolic genes from amplicon and metagenome libraries, respectively, from published microbial assessments of hydrocarbon reservoir fluids. By combining amplicon and metagenome surveys, this strategy unveils diverse taxonomic assemblages and ecosystem functional potential to identify global patterns in petroleum reservoir microbiomes.

## Methods

### Data acquisition

To determine the community composition and functional potential of the petroleum reservoir microbiome, published studies with microbial assessments were selected based on the following criteria: (i) samples originating from a petroleum reservoir, (ii) sample collection points at the well head or directly associated infrastructure, and (iii) publicly available sequence data. Metadata including sample collection point and type, reservoir name, reservoir (and/or collection point) location, temperature and depth, and the use of primary or secondary (e.g., water or steam injection) production was compiled directly from the respective publications (***Table S1***). In instances where geographic coordinates of the reservoir were not provided, latitude and longitude have been approximated based on the location of the reported oil field.

### 16S rRNA gene sequence processing

Raw high throughput 16S rRNA gene amplicon sequence data was obtained from the National Center for Biotechnology Information’s (NCBI) Sequence Read Archive (SRA; Leinonen *et al*. 2011) by compiling sequence accession lists and implementing the *prefetch* and *fastq-dump* commands from the SRA Toolkit. The resulting amplicon dataset consists of 53 million raw sequence reads comprising 62 Gb of sequence information. Initial sequence processing was performed using the open source VSEARCH version 2.11.1 (Rognes *et al*. 2016). In instances where paired-end sequence data were available from the SRA, read pairs were merged based on a minimum read overlap length of 10 base pairs (bp) and a maximum permitted mismatch of 20% of the length of the overlap. Merged reads were filtered with a maximum expected error of 0.5 for all bases in the read, and minimum and maximum read lengths of 150 and 500 bp, respectively (informed by the reported amplicon size). Identical reads were dereplicated and annotated with their associated total abundance for each sample, prior to *de novo* chimera detection and removal using default parameters. Accounts of the reads retained at each processing stage are provided in ***Table S2***. The quality-controlled sequences are available in the figshare online open access repository at https://doi.org/10.6084/m9.figshare.c.5801735.v2.

The studies compiled in this meta-analysis were published over a period during which sequencing and accepted standards have been evolving. Despite applying quality control steps designed to include a larger range of studies and enable a broader global meta-analysis with maximum breadth, in some instances as few as 1.7% of the raw reads in a given library and 8.4% of raw reads in an entire study could be retained. This highlights how data quality can be an important and sometimes unresolved issue in microbiology investigations.

Quality-filtered 16S rRNA gene amplicon sequences from each high throughput library were concatenated into a single file for further processing. To account for the different target hypervariable regions chosen within studies and between studies, sequences with consistent taxonomic classification were considered members of the same phylotype, allowing all libraries to be compared regardless of PCR methodology. Taxonomy was assigned using Mothur version 1.41.3 (Schloss *et al*. 2009) and Ribosomal Database Project’s k-mer-based naïve Bayesian classifier (Wang *et al*. 2007) with the SILVA SSU Ref NR version 138 database (Quast *et al*. 2013). A bootstrap cut-off value was set to 80% to return only taxonomies above this confidence threshold. A custom R script written in base R version 3.6.1 (R Core Team, 2014) parsed the output taxonomic assignments into a sample-by-phylotype table that accounted for earlier dereplication of the sequences. Sequence processing included the removal of singleton phylotypes, phylotypes with no domain level taxonomic classification (likely artifacts of sequence preprocessing), and phylotypes classified as *Vertebrata, Mitochondria* and *Chloroplast*. High throughput sequencing libraries with <1,000 reads after quality filtering were removed from the dataset. To limit the loss of data, subsampling was only used for alpha diversity calculations and for comparisons of the effect of variable library sizes (results are reported alongside non-subsampled libraries in specific instances as described herein). Subsampling for alpha diversity was performed without replacement to 1,000 reads using the *phyloseq* R package (McMurdie and Holmes, 2013). For all other analyses, subsampling was not performed in order to maximize the depth of information extracted from amplicon sequencing libraries.

Several oil reservoir amplicon sequencing studies have employed clone libraries (e.g., Voordouw *et al*. 1996; Gittel *et al*. 2009; Hubert *et al*. 2012). Amplicon sequencing reads derived from cloning were downloaded from NCBI’s GenBank database (Sayers *et al*. 2019) and analysed separately from the high throughput 16S rRNA gene amplicon sequence data described above, in order to account for the varying sampling depth and data outputs. No sequence processing steps were implemented prior to the analysis of these ‘low throughput’ amplicon libraries, except for using the reverse complement sequence for taxonomy in instances where this produced a better classification. Analysed clone sequences are compiled in the figshare online open access repository at https://doi.org/10.6084/m9.figshare.c.5801735.v2. Taxonomy was assigned using Mothur version 1.41.3 (Schloss *et al*. 2009) and Ribosomal Database Project’s k-mer-based naïve Bayesian classifier (Wang *et al*. 2007) with the SILVA SSU Ref NR version 138 database (Quast *et al*. 2013). A bootstrap cut-off value was set to 80%. Consistent taxonomic classification meant phylotype-level community structures between high throughput and low throughput amplicon libraries were comparable. Counts of the respective classifications were used to assess prevalence across the dataset at various taxonomic resolutions.

### Metagenome processing

The metagenomic dataset comprised sequences from 25 metagenome libraries – 16 raw libraries accounting for 457 million raw sequence reads were downloaded from NCBI’s SRA (Leinonen *et al*. 2011), in addition to 9.6 million assembled reads from nine libraries downloaded from the Integrated Microbial Genomes & Microbiomes system (IMG/M; Chen *et al*. 2021) when raw sequence data was not publicly available (***Table S2***). In total 448 Gb of sequence information was compiled. Raw, unassembled reads were quality-controlled by trimming technical sequences (primers and adapters) and low-quality additional bases, and filtering artifacts (phiX), low-quality reads and contaminated reads using BBDuk (BBTools suite, http://jgi.doe.gov/data-and-tools/bbtools). Ribosomal rRNA genes in quality-controlled reads were reconstructed and classified using phyloFlash (Gruber-Vodicka *et al*. 2017) with mapping against the SILVA SSU Ref NR version 138 database (Quast *et al*. 2013). Ribosomal rRNA genes in assembled contigs were identified using rRNAFinder (https://github.com/xiaoli-dong/rRNAFinder) after contigs smaller than 500 bp were removed, and taxonomy assigned using Mothur version 1.41.3 (Schloss *et al*. 2009) with the SILVA SSU Ref NR version 138 database (Quast *et al*. 2013). Detailed accounts of the reads retained during processing and assembly statistics are provided in ***Table S2***. Known challenges associated with 16S rRNA gene assembly from metagenomes (Yuan *et al*. 2015) likely account for the lower gene counts in prior assembled data as compared to unassembled data. For the quality-controlled reads not already assembled, the metagenome libraries were assembled using MEGAHIT version 1.2.2 (Li *et al*. 2015) with default parameters, and contigs <500 bp removed.

Gene-centric metagenomic analysis assessed the functional potential of the community as a whole. By focusing on genes rather than assembling whole provisional genomes much larger proportions of the metagenomic libraries are retained for analysis. Protein coding genes in each assembly were predicted using Prodigal version 2.6.3 (Hyatt *et al*. 2010). Predicted gene sequences were compiled and made available in the figshare online open access repository at https://doi.org/10.6084/m9.figshare.c.5801735.v2. Predicted genes were annotated with Kyoto Encyclopedia of Genes and Genomes (KEGG) Orthology (KO) using GhostKOALA (Kanehisa *et al*. 2016). Metabolic pathways from KO assignments were reconstructed using KEGG Decoder (Graham *et al*. 2018). Sporulation genes were identified by manual searching for KO numbers of specific marker genes. Genes involved in aerobic and anaerobic activation of hydrocarbon compounds (i.e., indicators of hydrocarbon biodegradation capability) were annotated using the CANT-HYD database of phylogeny-derived hidden Markov models (Khot *et al*. 2021). The CANT-HYD trusted cut-off domain score was used to annotate hydrocarbon activation functions reported in ***Table S7*** and ***Figure 4***. For the identification of *assA*-like pyruvate formate lyase (*pflD*) genes, no domain-level score cut-off was used, and the highest scoring sequence for each of the respective metagenomes was retained for phylogenetic analysis (***Figure 5***). Sequences sharing homology with *assA*-like pyruvate formate lyases (*pflD*) in metagenomes, together with experimentally verified and hypothetical *assA*, *bssA*, and *nmsA* genes, were aligned using Clustal Omega (Sievers *et al*. 2011) before inference of a maximum likelihood phylogenetic tree using FastTree (Price *et al*. 2009). Phylogenetic trees were annotated using iTOL version 5.5 (Letunic and Bork, 2019).

### Statistical analysis and data visualization

Statistical analyses and visualization were performed using base R version 3.6.1 (R Core Team, 2017) or the specific R packages as indicated below. Richness and alpha diversity metrics were calculated for subsampled high throughput 16S rRNA gene sequence libraries using *phyloseq* (McMurdie and Holmes, 2013). Spearman’s rank correlation coefficient assessed correlations between alpha diversity indices and two-group variables. Kruskal-Wallis tests assessed alpha diversity variance with respect to multi-group and non-numeric variables.

Prior to beta diversity calculations the non-subsampled and subsampled high throughput 16S rRNA gene amplicon datasets were split according to the implied and probable use of universal or archaeal primers; libraries comprising ≥75% archaea (based on relative sequence abundance) are defined here as archaeal libraries and the remainder defined as universal libraries indicating non-taxon specific 16S rRNA gene amplification. Non-metric multidimensional scaling (NMDS) of Bray-Curtis dissimilarity between high throughput 16S rRNA gene-based communities was calculated from relative abundance data using *phyloseq* (McMurdie and Holmes, 2013) and visualized using *ggplot2* (Wickham and Chang, 2015). Statistical differences between sample community dissimilarities (Bray-Curtis) in relation to non-numeric variables were assessed using Permutational Multivariate Analysis of Variance (PERMANOVA) tests in *vegan* (Oksanen *et al*. 2016). Biogeographical patterns were assessed using Mantel tests of sample community dissimilarities (Bray-Curtis) and the Euclidean distance between environmental parameters or haversine (geographic) distance between locations. Haversine distance was calculated using the *geosphere* package (Hijmans, 2019) and Mantel tests performed using *vegan* (Oksanen *et al*. 2016).

Microbial indicator species analysis, designed to test the association of a single taxon with an environment through multilevel pattern analysis, was used to identify phylotypes that best represent a specific environmental condition based on both phylotype presence/absence and relative abundance patterns. Indicator phylotypes were calculated using the *multipatt* function of the *indicspecies* package in R, employing a point-biserial correlation index (De Cáceres *et al*. 2010). Tests were performed on non-subsampled libraries (see ***Table S1*** for temperature subsets).

## Results

### Overview of the petroleum reservoir microbiome

A total of 2,473 phylotypes were inferred from high throughput 16S rRNA gene amplicon sequencing (***Table S5***) of 295 subsurface oil reservoir samples. Phylotypes encompassed 96 phyla comprising 2,283 genera in total. At the phylum level, *Proteobacteria* (30%), *Euryarchaeota* (19%), *Halobacterota* (15%), *Firmicutes* (9%) and *Campylobacteria* (5%) were most abundant. Classes and genera present at >1% average relative sequence abundance all belong to the ten most abundant phyla in the dataset (***Figure 1***). At the genus level, *Thermococcus, Methanosaeta*, and *Methanothermobacter* cumulatively account for 8–4% of the sequences. Oil reservoir clone libraries (48 samples comprising 2,850 reads) support findings from the more extensive high throughput 16S rRNA gene data, with *Halobacterota*, *Proteobacteria* and *Firmicutes* accounting for 26, 20 and 13%, respectively, at the phylum level. Oil reservoir metagenomes corroborate 16S rRNA gene sequencing approaches in terms of the most prevalent taxonomic groups, but highlight differences in relative abundance estimates (***Figure 1***). The most abundant phyla in metagenome libraries were *Firmicutes* (27%), *Proteobacteria* (20%), *Euryarchaeota* (9%), *Desulfobacterota* (8%), and *Thermotogota* (6%), corresponding with the most abundant genera in the metagenomes being *Caminicella*, *Pseudomonas*, *Desulfonauticus*, *Petrotoga*, and *Thermoanaerobacter*, each accounting for 15–4% of the genus-level assignments. Inconsistencies in community composition using these different DNA sequencing strategies may reflect true variations between the reservoirs sampled (N.B. few reservoirs in this study had both amplicon and metagenome sequencing applied), or could reflect well-known methodological differences such as preferential amplification of certain phylotypes during 16S rRNA gene sequence library preparation skewing the reported community composition.

**Figure 1.**
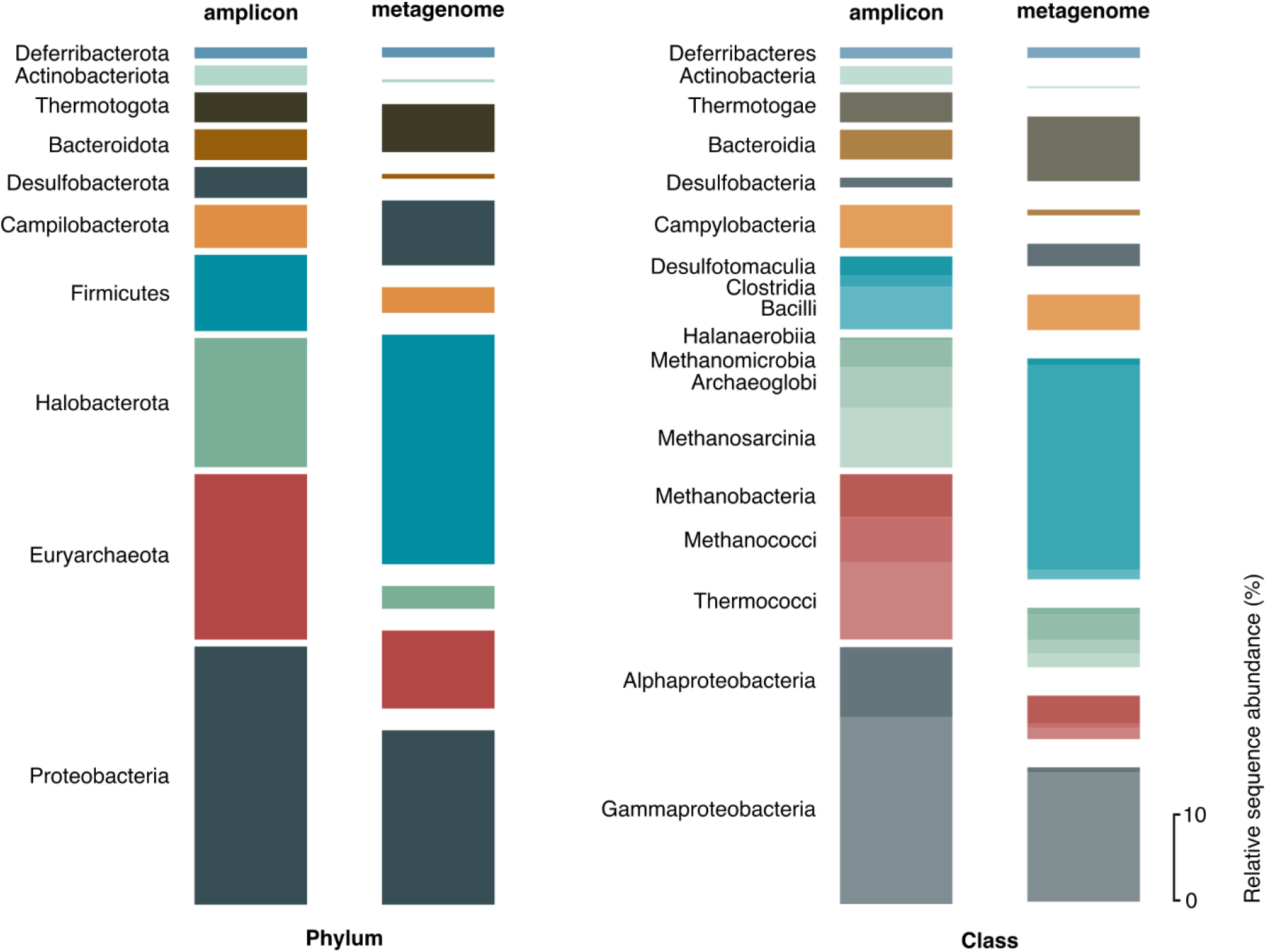
Petroleum reservoir microbiome. Average relative sequence abundance of the ten most abundant phyla and all classes >1% relative sequence abundance in 295 high throughput 16S rRNA gene amplicon sequencing libraries. The corresponding abundance of the same phyla and classes in 25 metagenome libraries are also shown. Phyla represent 93% and 78% of the 16S and metagenomes libraries, respectively. Classes represent 90% and 70%, respectively.

Assessing whether or not petroleum reservoirs around the world harbour a core microbiome revealed that the bacterial phyla *Proteobacteria* and *Firmicutes* and the archaeal phylum *Halobacterota* were observed across >75% of the high throughput 16S rRNA gene libraries. Accordingly, the classes *Gammaproteobacteria*, *Alphaproteobacteria*, *Methanosarcinia*, *Methanomicrobia, Clostridia* and *Bacilli* were identified in 82–60% of the samples. In metagenomic datasets, *Firmicutes* and *Proteobacteria*, along with *Euryarchaeota*, were observed in all samples, and at the genus level *Thermovirga, Halanaerobium* and *Petrotoga* were observed in more than 75% of the samples. Considering oil reservoir samples associated with primary recovery (*n* = 98), all high throughput 16S rRNA libraries showed Archaea, while *Firmicutes* and *Proteobacteria* were detected in 97 out of 98 libraries, and likely represent the best candidates for a core microbiome. Comparative analysis of reservoirs produced by secondary recovery methods showed *Proteobacteria* and *Halobacterota* were the most prevalent phyla, detected in 75 and 69% of the 198 libraries, respectively.

### Environmental influences on the community structure within oil reservoirs

Depth and, by association, temperature of the 368 petroleum reservoir samples spanning six continents ranged from 270–3,550 meters below surface (mbsf; defined as the seabed in offshore settings) and 8–110°C (***Table S1***). Alpha diversity and depth are negatively correlated in high throughput amplicon libraries (Spearman’s rank correlation *r* = −0.47 and −0.49, respectively, *P* < 0.05), with the deepest and hottest samples (>2,000 m and 74°C average reservoir temperature; *n* = 81; ***Table S1***) having a Shannon diversity index *H’* of 1.6 on average, compared to an average *H’* of 2.8 in the shallowest and coolest reservoirs (<500 m and 12°C average reservoir temperature; *n* = 76) (***Table S2***). This correlation of temperature with microbial diversity is apparent for oil reservoirs situated both onshore (*n* = 231) and offshore (*n* = 64).

Reservoir temperature and depth, as with alpha diversity, are correlated with greater variation in high throughput 16S rRNA gene-based community composition, such that as the difference in temperature and depth of two compared samples increases, community composition variance increases (***Figure 2**; **Table S4***). Assessments of phylotype presence and abundance for distinct reservoir temperature intervals using *indicspecies* tests (De Cáceres *et al*. 2010; ***Table S6***) showed that most members within the commonly observed phyla (e.g., *Proteobacteria, Halobacterota* and *Firmicutes*) were not significantly associated with an individual temperature interval (i.e., ten-degree intervals spanning 0–90°C). By contrast, at the genus level, *Thermococcus* and *Petrotoga* demonstrated strong correlations with temperature, being prevalent community members in reservoirs between 71–80 and 81–90°C, respectively. This was consistent with results from metagenomes, particularly for *Petrotoga* (***Table S5***).

**Figure 2.**
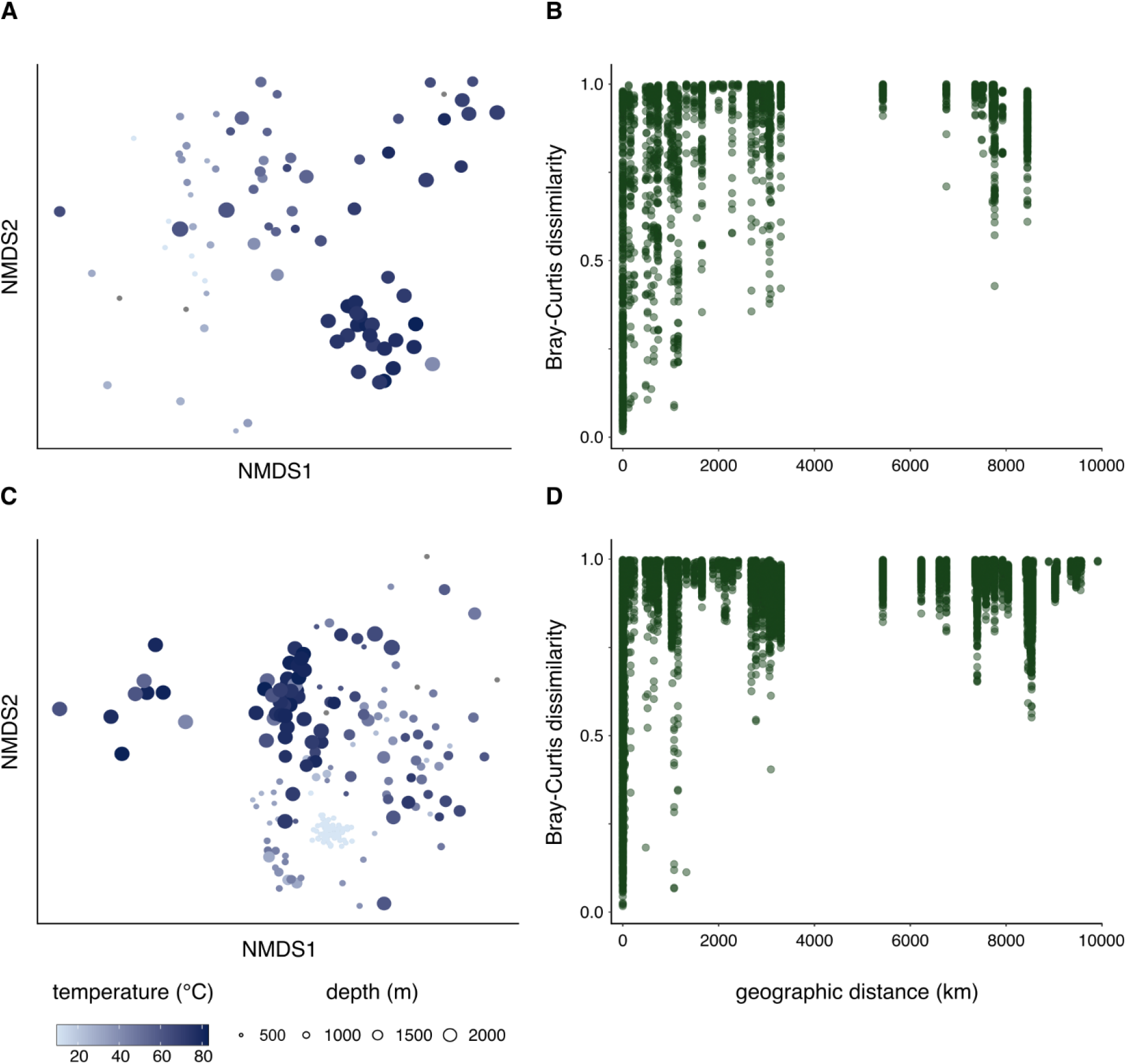
Relationship between microbial community composition and environmental factors. High throughput 16S rRNA gene libraries were grouped based on the use of Archaea-specific primers (**A**, **B**) or universal primers (**C**, **D**) to perform Bray-Curtis community dissimilarity comparisons based on temperature, depth and geographic distance. Libraries prepared using archaeal primers (**A**) exhibit greater dissimilarity between samples as a function of temperature (Mantel, *r* = 0.55) and depth (Mantel, *r* = 0.55) than libraries prepared with universal primers (**C**) (Mantel, *r* = 0.33 and 0.43, respectively), although both represent significant dissimilarity (*P* <0.05). Visual trends as well as statistical testing (Mantel, *P* < 0.05) in amplicon libraries using archaeal (**B**) and universal (**D**) primers indicate that geographically distant reservoirs have more dissimilar microbial communities.

Oil reservoirs undergoing secondary recovery operations (*n* = 197) are less diverse (average *H’* = 1.7) than reservoirs sampled during primary recovery (*n* = 98; *H’* = 2.8). Whereas primary oil recovery is mainly driven by *in situ* reservoir pressure, secondary oil recovery involves re-establishing pressure in the reservoir after the initial primary production phase by injecting fluids (e.g., seawater). Opportunities for exogenous organisms to be introduced into reservoir microbiomes during secondary recovery using non-sterile fluids could be expected to result in higher diversity as measured by the Shannon index and other metrics. The data here reveal the opposite, consistent with secondary recovery stimulating enrichment of opportunistic populations within the microbiome as environmental conditions change.

Primary or secondary oil recovery methods accounted for only a small fraction of the variation in microbial community composition (***Table S4***). Correlations were also apparent between geographic location and reservoir microbiome similarity (***Figure 2B***). As such, reservoirs in similar locations tend to have more similar microbial community compositions than reservoirs separated by large geographic distances. This distance decay relationship is more prominent if only reservoirs produced by primary recovery are considered (Mantel, archaeal *r* = 0.87, universal *r* = 0.80, *P* < 0.05) as compared to when reservoirs produced by secondary recovery are incorporated into geographic distance analysis (Mantel, archaeal *r* = 0.44, universal *r* = 0.30, *P* < 0.05). This suggests a biogeography especially in pristine oil reservoir deep biosphere environments that is diminished after populations shift in response to secondary recovery. Re-pressurization via fluid injection into subsurface oil reservoirs during secondary recovery may thus exert a normalizing effect on the microbiome regardless of geographic location. This is not necessarily because exogenous fluids contain similar organisms but rather could be due to the fluid chemistry provoking a re-organization of microbial populations already in the reservoir, e.g., introducing oxidants into a highly reduced biogeochemical system stimulates certain populations while limiting others.

### Functional potential of the oil reservoir microbiome

Metagenomes from 25 samples associated with both primary and secondary recovery, and from reservoirs ranging from 457–3350 mbsf and 26–102°C were compiled for community-wide gene prediction to assess the functional potential of the oil reservoir microbiome. In total 448 Gb of information was evaluated. Analysis of 6,992 unique KEGG Orthology (KO) assignments shows the functional composition of communities is highly conserved across samples from different studies (***Figure 3A**; **Table S4***). Unlike 16S rRNA taxonomy-based associations of microbiomes with environmental factors (***Figure 2, 3B***), no significant correlation was observed between gene composition and reservoir type (i.e., primary vs secondary recovery), temperature, depth or geographic location (***Table S4***). This is consistent with a functional core set of genes in subsurface oil reservoirs.

**Figure 3.**
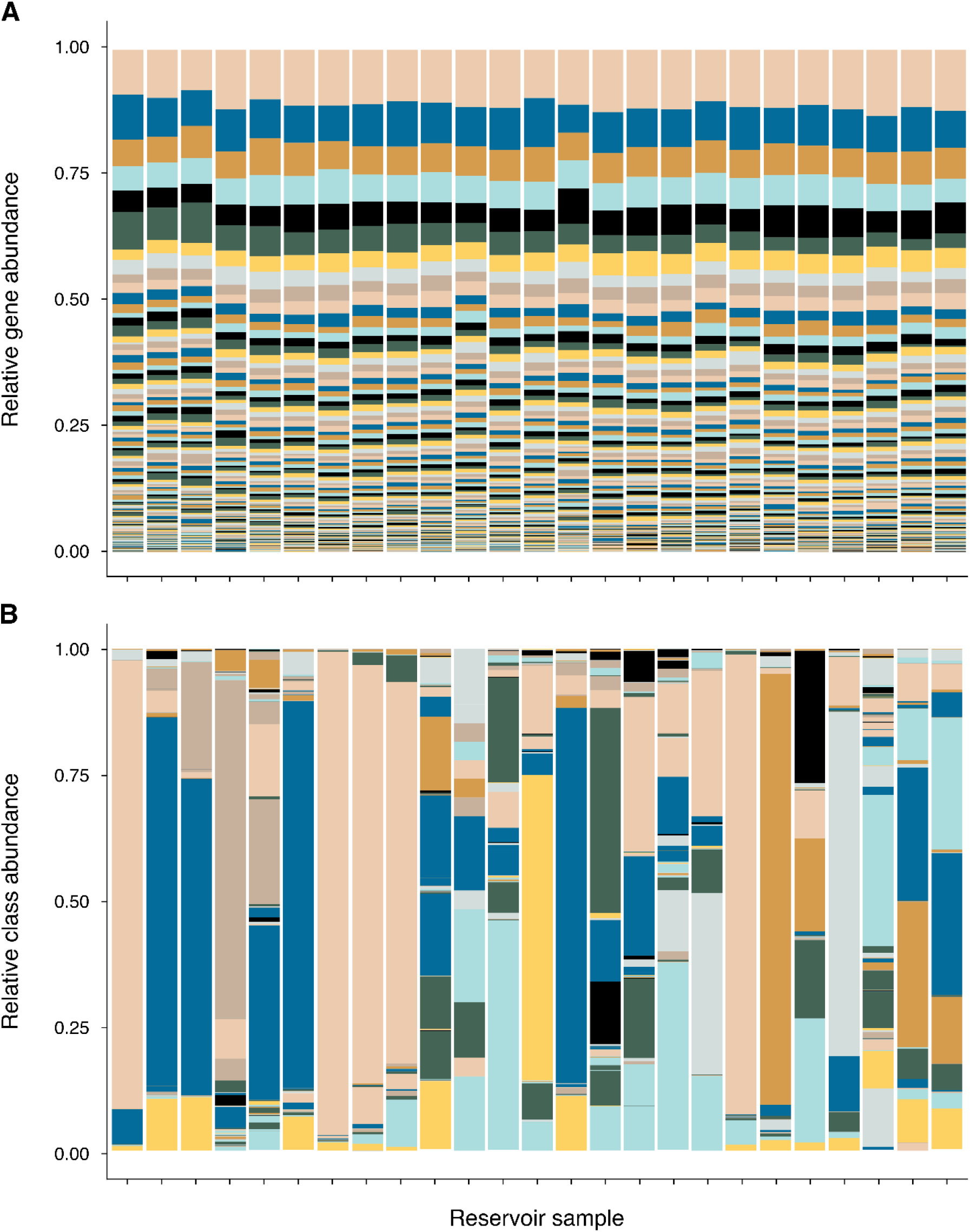
Metabolic gene and taxonomic composition of samples from different reservoirs. **A**, Relative abundance of genes (represented by unique KEGG orthologs and grouped by KEGG level C pathway) in metagenome sequence libraries from 25 different petroleum reservoir samples. Ortholog counts were subsampled to the lowest total number of KEGG-annotated sequences across all samples in the dataset. **B**, Relative abundance of taxonomic groups (class level) in the same 25 metagenome sequence libraries. The similar gene composition profiles and variable taxonomy-based associations reflect a conserved genetic potential of the petroleum reservoir microbiome regardless of prevailing taxa in these habitats (***Table S4***).

Reservoir metagenomes were compared to better understand functional potential for carbon acquisition and energy conservation (***Figure 4***; ***Table S7***). Diverse capabilities for carbohydrate, peptide, and lipid catabolism, as well as carbon fixation, are widespread in oil reservoirs. Mixed-acid fermentation appears to be a universal strategy. Acetate production from pyruvate is observed in all metagenomes, consistent with acetate often being measured in oil field waters (Carothers and Kharaka, 1978; Barth, 1991) and acetogenesis being important in the deep biosphere in general (Lever, 2012). Energy conservation through anaerobic respiration is another widespread metabolism. Genes required for the reduction of sulfate to sulfide (*sat*, *aprAB* and *dsrAB*) were observed in 24 of the 25 metagenomes. Reductases required for nitrate reduction to nitrite, dissimilatory nitrate reduction to ammonia (DNRA) or denitification to N_2_ were detected in 18, 15, and 8 of the metagenomes, respectively. Observations of sulfate and nitrate metabolism genes do not correlate with primary or secondary recovery practices, indicating that these respiratory pathways are universal features of indigenous microbial communities. Sulfide:quinone oxidoreductase (*sqr*), which catalyzes the oxidation of sulfide to elemental sulfur, was detected in all samples. Sulfide oxidation can be coupled to DNRA or denitrification (Hubert and Voordouw, 2007), which these oil reservoir microbiomes generally exhibit capability for, as noted above. Biological oxidation represents a strategy for mitigating the accumulation of sulfide (Marcia *et al*. 2009) that is generated in oil reservoirs as a result of biological or thermochemical sulfate reduction (Machel *et al*. 1995).

**Figure 4.**
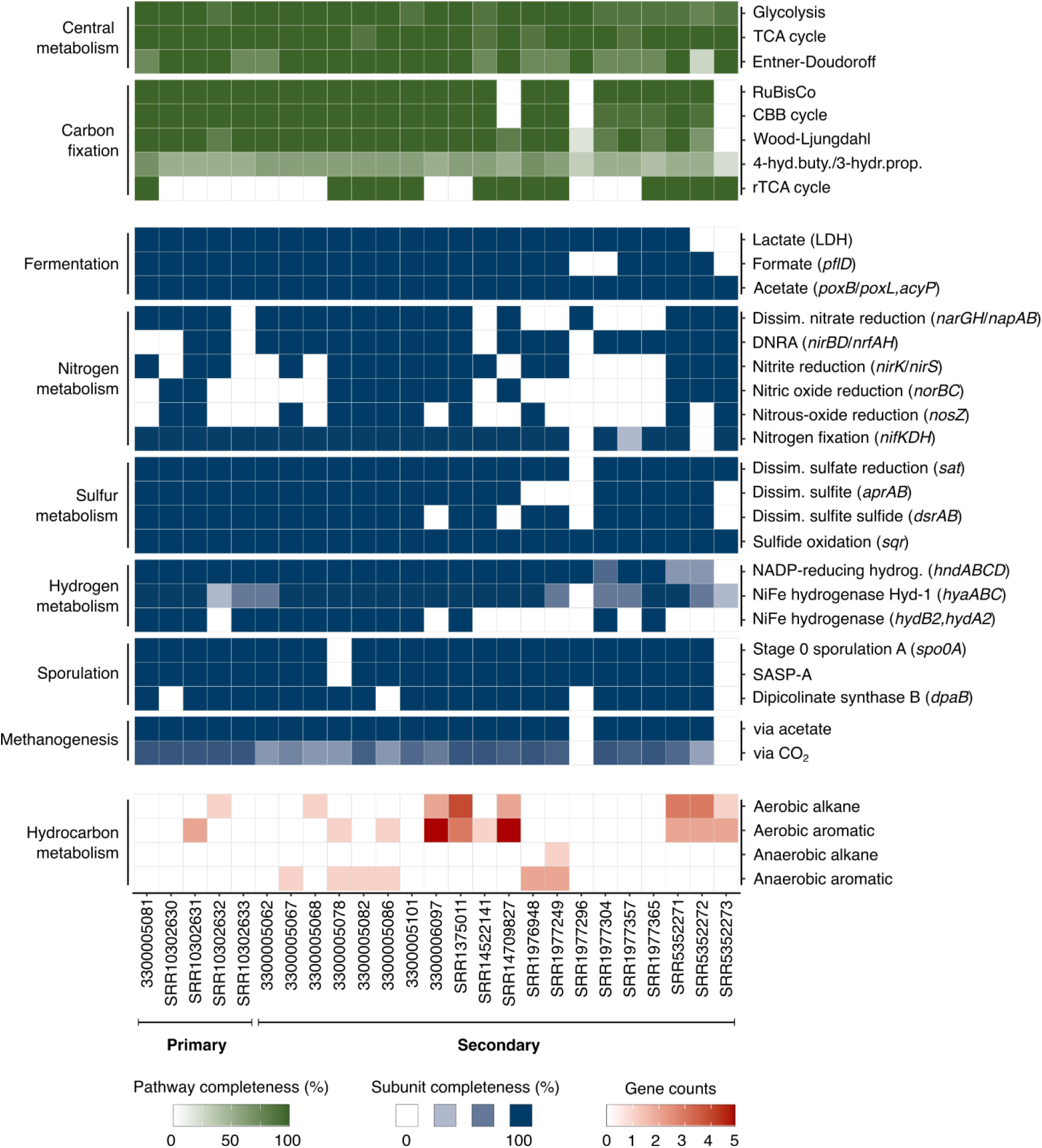
Metabolic potential of the petroleum reservoir microbiome. Occurrence of metabolic genes and pathways in 25 metagenomes from microbiomes from primary and secondary produced reservoirs. Complete gene lists can be found in ***Table S7***.

Hydrogen metabolism in oil reservoirs can be linked to both fermentative and respiratory pathways (***Figure 4***; ***Table S7***). Membrane-bound Group 1b respiratory H_2_-uptake hydrogenases (*hydB2, hydA2*) and Group 1d oxygen-tolerant NiFe hydrogenase (*hyaABC*) are both observed in the metagenomes. These enzymes pair hydrogen oxidation with the liberation of electrons for anaerobic respiration (Greening *et al*. 2016). Consistent co-occurrence of *hyaABC* with the near universal dissimilatory sulfate reduction gene sulfate adenylyltransferase (*sat*) suggests potential coupling of this form of hydrogen oxidation with sulfate respiration, if sulfate is present. Genes encoding the Group 3b cytosolic NiFe hydrogenase (*hndABCD*) were present in nearly all metagenomes. This enzyme can work bidirectionally, but generally couples reoxidation of NAD(P)H to H_2_ generation via fermentation (Greening *et al*. 2016). The diversity of hydrogenases in oil reservoir metagenomes suggests that both hydrogen production and consumption may be widespread features of oil reservoir biogeochemistry.

Microbial dormancy in the form of endospore formation is considered to be an important process in the deep biosphere (Wörmer *et al*. 2019). Key marker genes for sporulation are widespread in these subsurface oil reservoir metagenomes (***Figure 4***; ***Table S7***). These include the *spo0A* master transcriptional response regulator that controls entry into sporulation, and genes encoding small acid-soluble protein (SASP-A) and dipicolinic acid (*dpaB*) synthesis that are important in protecting spore DNA against damage. This finding is consistent with the *Firmicutes* phylum (which known endospore-forming microbes belong to) demonstrating the highest average relative abundance in oil reservoir 16S rRNA gene and metagenome libraries (***Figure 1***).

### Hydrocarbon biodegradation potential

To identify potential for microbial degradation of crude oil, functional marker genes encoding enzymes that initiate aerobic or anaerobic hydrocarbon biodegradation, by activating either alkane or aromatic compounds, were examined using a set of 37 newly developed hidden Markov models (Khot *et al*. 2021; ***Figure 4***; ***Table S7***). Of the 28 aerobic and 9 anaerobic marker genes tested, 17 were identified in 16 of the 25 metagenomes. Among these 17 genes, 14 are associated with the aerobic biodegradation of hydrocarbons. Genetic potential for the aerobic oxidation of *n*-alkanes with chain lengths C_10_-C_16_ by *alkB* alkane hydroxylase and short- and medium-chain-length *n*-alkanes by the cytochrome P450 CYP153 family alkane hydroxylase was prevalent. Genes involved in the aerobic degradation of long chain alkanes (e.g., *almA* and *ladA/B*) and polyaromatic hydrocarbons (e.g., *ndoB/C* and *dszC*) were less frequently observed.

Anaerobic degradation of *n*-alkanes and aromatic hydrocarbons can be initiated by their addition to fumarate to form succinates. Sequences encoding the catalytic subunit of benzylsuccinate synthase (*bssA*) were detected in six samples from reservoirs in the North Sea and North Slope Alaska, with the latter also hosting the catalytic subunits of alkylsuccinate synthase (*assA*) and naphthylmethylsuccinate synthase (*nmsA*). Metabolisms encoded by these genes can couple hydrocarbon degradation to the reduction of an electron acceptor such as sulfate, or to fermentative or syntrophic metabolisms that proceed in conjunction with methanogenesis — a process likely to be a primary mechanism for hydrocarbon metabolism in reservoirs worldwide over geological timescales (Jones *et al*. 2008; Gray *et al*. 2011). Genes involved in acetoclastic and hydrogenotrophic methanogenesis that facilitate this were detected in 23 of the 25 metagenomes. Similarly, the gene encoding the Group 3a: F420 hydrogenase, *frhB* (KEGG ortholog KO00441), which directly couples oxidation of hydrogen to the reduction of F420 during methanogenesis (Mills *et al*. 2013) is widespread, being detected in 23 of the 25 metagenomes (***Table S7***).

## Discussion

### Petroleum reservoirs lack a core microbiome

By compiling 295 amplicon libraries from 41 oil reservoirs totalling 53 million sequence reads, this study was able to falsify the hypothesis that oil reservoirs contain a core microbiome, defined as the presence of common species in all samples (Shade and Handelsman, 2011). Taxonomic assessments showed that even at the phylum level not all of the samples included representatives of the most prevalent groups, i.e., *Firmicutes, Proteobacteria* and *Halobacterota*. Interestingly, the proportion of samples with these phyla was higher if only reservoirs produced by primary oil recovery are considered, but even then, these groups were not universal. Taxonomic assessment of contigs from metagenomic libraries in a smaller number of samples supported observations from amplicon libraries indicating that no phyla are universally present in oil reservoirs.

Instead of a core microbiome in oil reservoirs, environmental factors result in niche habitat partitioning whereby taxonomically different organisms comprise the microbiome in oil reservoir habitats featuring different physicochemical conditions. Metagenomic sequencing libraries from 25 reservoir samples reveal that different oil reservoirs share a common functional core (***Figure 3A; Table S4***). Redundant functional potential in different oil reservoirs demonstrates trait selection in these settings despite the absence of a taxonomic core (***Figure 3B***). A diversity of genes encoding functions for relevant processes such as fermentation, sulfate reduction, hydrocarbon biodegradation and methanogenesis is indicative of metabolic versatility being an important feature of the oil reservoir microbiome. These and other metabolic functions were highlighted in a similar meta-analysis of oil reservoir metagenomes by Hidalgo *et al*. (2021), which assessed metagenome-assembled genomes (MAGs) from nine different oil reservoirs in China, Alaska’s North Slope and offshore Brazil. That genome-centric approach revealed both core and environment-specific functions, but also included observations that differ from those in the genecentric analysis presented here with respect to the metabolisms noted above. Some inconsistency between these two metagenome analysis strategies may be expected to arise based on preferential assembly of the most abundant organisms in genome-centric analyses resulting in an associated loss of unbinned sequence information in the process. For example, MAGs containing genes for dissimilatory sulfate reduction were identified in just over half of the oil fields in the genome-centric study (Hidalgo *et al*. 2021) whereas in the analysis performed here, without a genome binning step, sulfate reduction genes were identified in 24 out of 25 samples. Similarly, genome-centric analysis suggested that sulfide oxidation was not widespread in oil fields, whereas the less restrictive analysis of unassembled contigs in the present study suggests a universal genetic potential for sulfide oxidation in oil reservoirs. This highlights the different ways that metagenomic surveys can be assessed in terms of delivering greater understanding of the metabolic potential of oil reservoir microbial communities.

Genetic capacity for carbohydrate, peptide, and lipid metabolism, uncovered in the present study, indicates that degradation of microbial necromass and sedimentary detrital material could be an important process in petroleum reservoirs, or their precursor sediments. This is consistent with the role of residual organic matter recycling in the deep biosphere in general (Lomstein *et al*. 2012; Lloyd *et al*. 2013; Orsi *et al*. 2013). The widespread occurrence of genes involved in acetogenesis in these metagenomes is also consistent with acetogens being important in anaerobic subsurface environments (Lever, 2012). Diversity and prevalence of both respiratory and fermentative pathways indicates these may be co-occurring or used sequentially by individual organisms in response to changing conditions. Whereas respiration has favourable energetics relative to fermentation, the limited availability of electron acceptors in the subsurface is likely an important determinant for the predominance of fermentation, which may additionally reflect a strategy to access energy from more challenging substrates such as hydrocarbons. It is likely that fermentation products contribute substrates for further sulfate reduction or methanogenesis, depending on the prevailing redox conditions. Not surprisingly, genes for sulfate reduction and methanogenesis were widespread in the 25 oil reservoir metagenomes examined here. The potential for respiration may also explain the rapid transition to souring and souring control scenarios following the introduction of sulfate-rich seawater and injected nitrate, respectively.

### Is hydrocarbon biodegradation universal in petroleum reservoirs?

In pristine oil reservoirs, prior to secondary recovery or even reservoir discovery, the absence of oxygen or other electron acceptors dictates that methanogenic conditions should prevail in many settings. In this context, fermentation reactions that initiate anaerobic hydrocarbon biodegradation couple with methanogenic archaea rapidly consuming acetate, CO_2_ and hydrogen to ensure thermodynamic feasibility of syntrophic partnerships (Schink, 1997; Dolfing *et al*. 2008). Recent work demonstrates that similar metabolism (conversion of oil to methane) can also be facilitated by oil reservoir methanogens in the absence of a syntrophic partner (Zhou *et al*. 2021). Genetic potential for acetoclastic and hydrogenotrophic methanogenesis was identified in all of the oil reservoirs examined here, consistent with the understanding that this may be a default biogeochemical regime in pristine petroleum-bearing sediments — a view supported by thermodynamic modelling (Dolfing *et al*. 2008), carbon dioxide and methane stable carbon isotopic measurements (Jones *et al*. 2008), and radiotracer experiments (Mayumi *et al*. 2011) that predict hydrogenotrophic CO_2_ reduction to be the primary route for crude oil hydrocarbon biodegradation in oil reservoirs.

On the basis of this evidence for anaerobic degradation of hydrocarbons being prevalent in these environments, corresponding hydrocarbon activation genes are expected to be widespread. However, *assA*, *bssA* and *nmsA* were only detected in two of the eight reservoirs examined by metagenome sequencing here. A similar result was reported by Hidalgo *et al*. (2021) who found only two *bssA* genes in 148 MAGs from different oil reservoirs using KEGG assignments. The discrepancy of widespread genomic potential for methanogenesis and geochemical evidence of biogenic methane production in reservoirs (Horstad and Larter, 1997; Larter and di Primio, 2005) on the one hand, and limited occurrences of anaerobic hydrocarbon degradation genes on the other, hints that other mechanisms for anaerobic hydrocarbon biodegradation (e.g., Lack and Fuchs, 1994; Heider *et al*. 2016; Rabus *et al*. 2016) or as-yet unknown forms of hydrocarbon activation in anoxic petroleum reservoirs. For example, analysis of variants of glycyl-radical enzymes proposed to mediate anaerobic alkane biodegradation via addition to fumarate reveals a clade of genes encoding alkylsuccinate synthase (*assA*) that is divergent from canonical *assA* found in *Proteobacteria* (***Figure 5***). These *assA*-like pyruvate formate-lyase (*pflD*) genes are found in taxa from different petroleum reservoirs, including *Archaeoglobus fulgidus* (Birkeland *et al*. 2017; Khelifi *et al*. 2014), ^U^*Petromonas tenebris* (Christman *et al*. 2020) and *Thermococcus sibiricus* (Mardanov *et al*. 2009). In-depth analysis of the predicted genes in each of the reservoir metagenomes assessed here shows that 16 out of the 25 libraries contained homologs of *assA* -like pyruvate formate-lyase (*pflD*) genes (***Figure 5***), pointing to underexplored potential for anaerobic hydrocarbon biodegradation within the petroleum reservoir microbiome.

**Figure 5.**
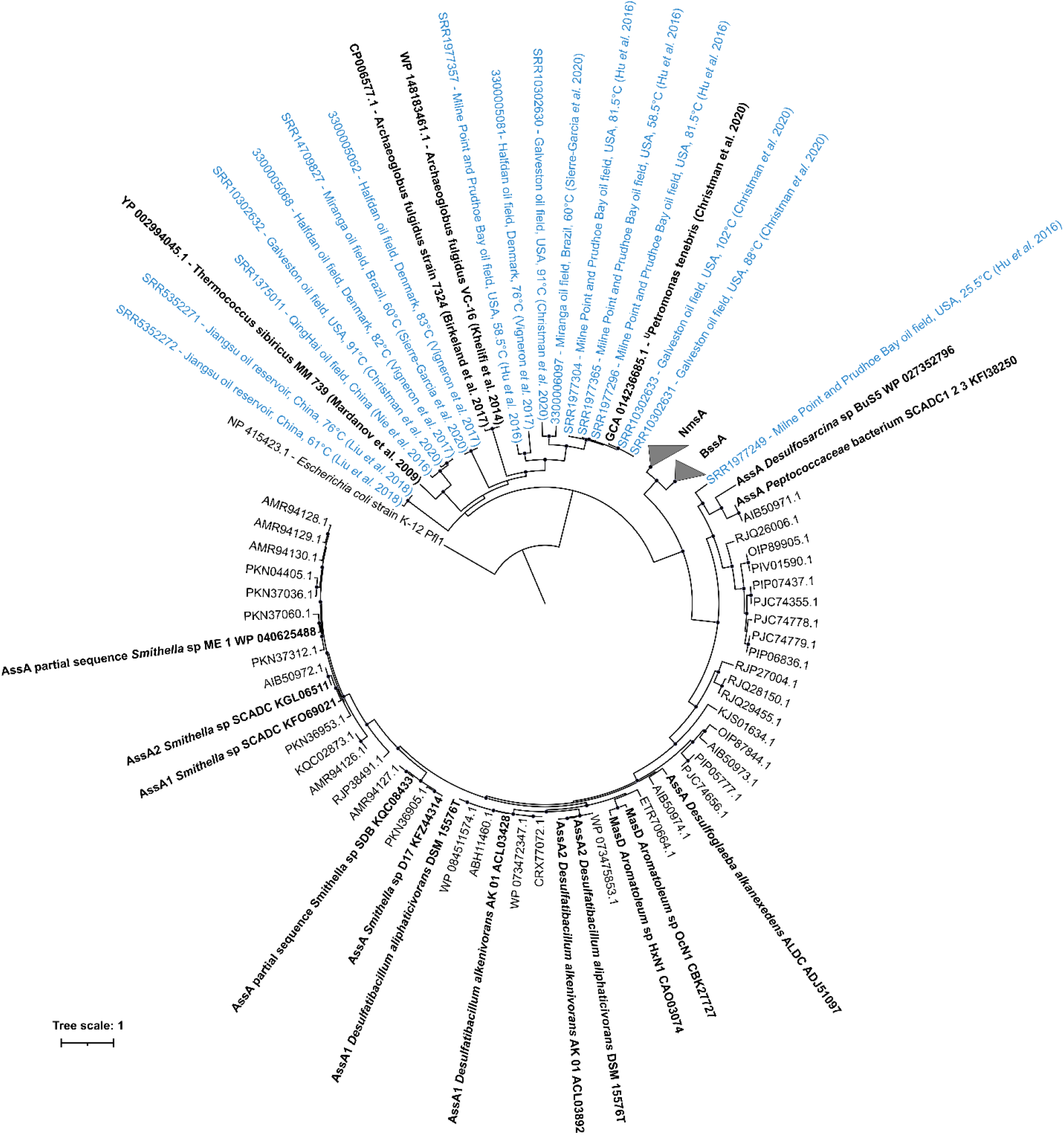
Phylogeny of *assA*-like pyruvate formate-lyase (*pflD*) genes. Putative anaerobic alkane-activating pyruvate-formate lyase enzyme variants from 16 of the 25 metagenomes analysed in this study (shown in blue) cluster together with homologous *pflD* gene sequences found in oil reservoir thermophiles ^U^*Petromonas tenebris* (Christman *et al*. 2020), *Archaeoglobus fulgidus* strain 7324 (Khelifi *et al*. 2014; Birkeland *et al*. 2017), and *Thermococcus sibiricus* strain MM 739 (Mardanov *et al*. 2009). *Archaeoglobus* and *Thermococcus* strains have been shown to degrade alkanes in pure culture (Khelifi *et al*. 2014; Mardanov *et al*. 2009). These pflD gene sequences, as well as other experimentally-verified alkane succinate synthase (AssA and MasD) genes, are shown in bold. Benzyl succinate synthase (BssA) and naphthyl-2-methyl-succinate synthase (NmsA) sequences are represented by collapsed clades. Black circles at the branch nodes indicate >80% bootstrap support (1,000 re-samplings). Scale bar indicates 10% sequence divergence as inferred from PhyML. The tree is rooted using Pfl from *E. coli*.

Detection of aerobic genes in many of the oil reservoir metagenomes could be considered surprising. Despite possible influx of oxygen-bearing meteoric waters (Palmer, 1993; Parkes and Maxwell, 1993), anoxic conditions are understood to prevail in subsurface petroleum reservoirs (Head *et al*. 2003). Interestingly, metagenomes containing aerobic genes were not restricted to samples associated with secondary oil recovery; long-chain alkane and monoaromatic dioxygenase enzymes are encoded in microbiomes from an offshore reservoir in the Gulf of Mexico (Christman *et al*. 2020) and an onshore reservoir in northeastern Brazil (Sierra-Garcia *et al*. 2020) that had not undergone secondary recovery prior to sampling for DNA analysis (***Figure 4***). These results are consistent with observations of high proportions of genes for enzymes involved in aerobic hydrocarbon metabolism in various anaerobic hydrocarbon resource environments including oil reservoirs (An *et al*. 2013; Hidalgo *et al*. 2021) and may arise from enrichment of initially minor groups of organisms capable of aerobic respiration coming into contact with air during sampling and sample transport. This is substantiated by genes encoding alkane hydroxylase (*alkB*) and monoaromatic dioxygenase (MAH) showing high sequence identity (based on BLAST against the non-redundant nucleotide database) with *Marinobacter* spp., which are generally detected in low relative abundance *in situ* by 16S rRNA gene amplicon sequencing (***Table S5***). Enrichment of aerobic bacteria *ex situ* during sampling is ecologically similar to proliferation of sulfate- or nitrate-reducing bacteria *in situ* in response to reservoir conditions changing, pointing more generally to an inactive or latent metabolic potential inherent to oil reservoir microbial communities.

### Provenance of the oil reservoir microbiome

The extensive nature of this global dataset of oil reservoir microbiomes suggests a pattern of subsurface biogeography featuring increasing community variance with greater geographic distance between reservoirs (***Figure 2***). This raises questions about the provenance of microbes and the establishment of the global reservoir microbiome in a geologic context. Heuer *et al*. (2020) postulated that vegetative cells and endospores that are deposited in surface sediments and undergo burial over geological timescales can then be revived under the right selective conditions. This is consistent with recent observations in a 1,180 m sediment core sampled during IODP Expedition 370 (Beulig *et al*. 2022), and the concept of a microbial dispersal loop proposed for understanding the interplay of ecological principles of selection and migration in the subsurface (Mestre and Höfer *et al*. 2020; Gittins *et al*. 2022). Aside from geological processes such as sedimentation and petroleum fluid migration, natural dispersal vectors in the deep biosphere are limited (Stetter *et al*. 1993). This evidence combines to suggest that microbial communities in oil reservoirs are inherited from populations that are present during the proximal deposition of sediments that eventually form reservoirs. Environmental selection during burial likely contributes to the differences observed here as a function of geography between oil reservoirs from different parts of the world.

While these principles may explain the assembly of microbial communities in pristine petroleum reservoirs, engineering interventions associated with oil production alter these environments dramatically. For example, reservoir re-pressurization in offshore settings results in introduction and mixing of seawater that typically features microbial diversity (Ladau *et al*. 2013; Zorz *et al*. 2019) that is greater than what was determined here in reservoirs that did not experience water injection (*H’* = 2.8). The number of reservoir samples assessed in this study enables an empirical assessment of the consequences of secondary oil recovery on the microbiome present in oil reservoirs, revealing that microbial diversity is actually lower in secondary produced fluids (*H’* = 1.7). It is well known that seawater injection is associated with oil reservoir souring. While the introduction of new organisms (Bell *et al*. 2020) or cooling of the near injection wellbore region to more permissive temperatures (Eden *et al*. 1993) are sometimes cited as factors that link water injection to souring, the large amount of sulfate present in seawater is understood to be a main driver for this outcome. This has led some oil companies to consider costly strategies for nanofiltration to remove sulfate from injected fluids (Bilstad, 1992). What is less clear is whether or not the sulfate-reducing microorganisms (SRM) responsible for souring are introduced exogenously via seawater injection or are persisting in the reservoir until conditions that favour this metabolic response arise. The former interpretation is challenged by seawater being an oxic environment where SRM may not be plentiful or active (Bell *et al*. 2020), whereas the latter would require SRM to persist in the reservoir in the absence of sulfate — a situation that may be explained by SRM that are also capable of fermentative metabolism (Muyzer and Stams, 2008). Similar questions can be asked regarding nitrate-reducing microorganisms (NRM) that are activated by nitrate injection for souring control, which they can achieve by coupling organotrophic metabolism to nitrate reduction to out-compete SRM, or by oxidizing sulfide directly with nitrate (Carlson and Hubert, 2019).

## Conclusion

DNA-based assessments of petroleum reservoirs continue to offer great potential for understanding difficult-to-access deep biosphere habitats and making important operational predictions in the face of changing environmental conditions. Designing and managing effective oil recovery strategies depend on the interplay between biogeochemical cycles catalyzed by microbiomes that are versatile and resilient to dramatic perturbations like seawater injection. The analyses presented here highlight the utility of 16S rRNA gene amplicon and shotgun metagenomic sequencing of oil reservoir samples. The extensive screening of hundreds of samples by amplicon sequencing demonstrates the effects of environmental selection and biogeography, and the absence of a core taxonomy, in oil reservoirs globally. Gene-centric metagenomic analysis reveals that this is possible thanks to a core functional biogeochemistry based on identifying widespread genes for key processes such as sulfate reduction, sulfide oxidation, nitrate reduction and methanogenesis. Less widespread detection of other genes, such as those suspected to catalyze anaerobic hydrocarbon biodegradation, raise important questions about these ecosystems, and suggest that microbiome investigations will continue to deliver insights into microbial processes of industrial and ecological relevance.

## Notes

https://doi.org/10.6084/m9.figshare.c.5801735.v2

### Competing Interest Statement

The authors have declared no competing interest.

